# Protection of Hamsters Challenged with SARS-CoV-2 after Two Doses of MVC-COV1901 Vaccine Followed by a Single Intranasal Booster with Nanoemulsion Adjuvanted S-2P Vaccine

**DOI:** 10.1101/2022.02.24.481901

**Authors:** Yi-Jiun Lin, Meei-Yun Lin, Ya-Shan Chuang, Luke Tzu-Chi Liu, Tsun-Yung Kuo, Charles Chen, Shyamala Ganesan, Ali Fattom, Vira Bitko, Chia-En Lien

**Affiliations:** Medigen Vaccine Biologics Corporation, Taipei City, Taiwan; Department of Biotechnology and Animal Science, National Ilan University, Yilan County, Taiwan; Temple University, Philadelphia, PA 19122, USA; BlueWillow Biologics, Ann Arbor, MI 48105 USA; Institute of Public Health, National Yang-Ming Chiao Tung University, Taipei City, Taiwan

## Abstract

Intramuscular vaccines have greatly reduced hospitalization and death due to severe COVID-19. However, most countries are experiencing a resurgence of infection driven predominantly by the Delta and Omicron variants of SARS-CoV-2. In response, booster dosing of COVID-19 vaccines has been implemented in many countries to address waning immunity and reduced protection against the variants. However, intramuscular boosting fails to elicit mucosal immunity and therefore does not solve the problem of persistent viral carriage and transmission, even in patients protected from severe disease. In this study, two doses of stabilized prefusion SARS-CoV-2 spike (S-2P)-based intramuscular vaccine adjuvanted with Alum/CpG1018, MVC-COV1901, were used as a primary vaccination series, followed by an intranasal booster vaccination with nanoemulsion (NE01)-adjuvanted S-2P vaccine in a hamster model to demonstrate immunogenicity and protection from viral challenge. Here we report that this vaccination regimen resulted not only in the induction of robust immunity and protection against weight loss and lung pathology following challenge with SARS-CoV-2, but also led to increased viral clearance from both upper and lower respiratory tracts. Our findings showed that intramuscular MVC-COV1901 vaccine followed by a booster with intranasal NE01-adjuvanted vaccine promotes protective immunity against both viral infection and disease, suggesting that this immunization protocol may offer a solution in addressing a significant, unmet medical need for both the COVID-19 and future pandemics.

## Introduction

The COVID-19 pandemic caused by SARS-CoV-2 (Severe Acute Respiratory Syndrome Coronavirus 2) is the worst pandemic the world has faced in over 100 years^1, 2^. Currently approved vaccines including (Pfizer-BioNTech BNT162b2 and Moderna mRNA-1273)^5, 6^ and adenovirus-based vaccines (Johnson & Johnson Ad26.COV2.S and Oxford/AstraZeneca AZD1222/ChAdOx1 nCoV-19)^7, 8^ have been highly effective at preventing severe disease and death in those vaccinated ^3, 4^. However, as the virus continues to mutate, novel variants of concern (VOC) such as the Delta and Omicron variants continue to result in waves of infection^9, 10^. In addition to mutant strains which may reduce vaccine efficacy, there is mounting evidence that the current vaccines lose potency over time^11^. Specifically, vaccinated people become more susceptible to infection starting about 6 months post vaccination^11^. Most importantly, intramuscular vaccination does not sufficiently prevent nasal shedding and transmission of the virus from person to person. Despite these challenges, intramuscular boosting remains the primary strategy for attempting to halt new waves of infection^12^.

COVID-19 infection occurs after virus-containing respiratory aerosols access the upper respiratory tract (URT) that includes the nasal cavity^13^. The nasal passage is the initial and most important route of infection due to the presence of large numbers of angiotensin-converting enzyme 2 (ACE2) and cellular serine protease TMPRSS2, both cellular proteins that are required for SARS-CoV-2 infection of nasal ciliated cells^14–16^. In addition, efficient viral replication of SARS-CoV-2 results in high viral titers in the nasopharynx which subsequently leads to lung infection and disease progression.

The main mode of SARS-CoV-2 transmission is through exposure to respiratory secretions or respiratory droplets, which are released from an infected person. Importantly, both symptomatic and asymptomatic infections have been confirmed to result in viral shedding ^16–19^. Consequently, the central role of nasal mucosa in both efficient SARS-CoV-2 infection and transmission has important implications for vaccine development. It has been shown that nasally administered vaccines that establish mucosal immunity at the port of viral entry and induce a systemic immune response are of considerable prophylactic value, as they can provide sterilizing immunity and block human to human transmission^20–22^. Intranasal vaccines already exist for indications other than infection by SARS-CoV-2 and have significant practical advantages over standard intramuscularly administered vaccines or orally administered antivirals^23^.

The current SARS-CoV-2 vaccines are almost all administered intramuscularly, and only offer partial protection against establishment of virus in the URT, which may explain the waves of SARS-CoV-2 infections observed since the initial COVID-19 outbreaks^24, 25^. Intranasal vaccines may address this issue via enhancement of localized nasal mucosal immunity and local immunological memory. This approach appears to be promising based on recent animal studies with a nasal spray version of ChAdOx nCoV-19 administered as two doses of primary vaccination or a Prime and Spike approach using mRNA intramuscular vaccine as a Prime followed by recombinant unadjuvanted spike protein by IN administration, as well as broad sarbecovirus mucosal immunity conferred by unadjuvanted recombinant spike protein delivered via mRNA-liponanoparticle^26, 27^.

MVC-COV1901 is a protein subunit vaccine based on stable prefusion SARS-CoV-2 spike protein S-2P adjuvanted with CpG 1018 and aluminum hydroxide^28^. MVC-COV1901 has been approved for use in Taiwan on the basis of its immunogenicity and safety profile^29, 30^. To develop an intranasal version of MVC-COV1901 using its core component S-2P, we carried out hamster studies to evaluate a vaccine of prefusion-stabilized S-2P formulated in NE01 adjuvant. We show that this vaccine is able to boost waning systemic immunity by increasing levels of anti-SARS-CoV-2 antibody after initial intramuscular vaccinations, in addition to inducing mucosal immune responses which protect hamsters from infection, viral carriage and disease following SARS-CoV2 challenge.

## Materials and Methods

### Animals and ethical statements

Female golden Syrian hamsters aged 8-10 weeks at study initiation were obtained from the National Laboratory Animal Center (Taipei, Taiwan). Animal immunizations were conducted in the Testing Facility for Biological Safety, TFBS Bioscience Inc., Taiwan. Three weeks after the final immunization, the animals were transferred to Academia Sinica, Taiwan, for SARS-CoV-2 challenge. All procedures in this study involving animals were conducted in a manner to avoid or minimize discomfort, distress, or pain to the animals and were carried out in compliance with the ARRIVE guidelines (https://arriveguidelines.org/). All animal work in the current study was reviewed and approved by the Institutional Animal Care and Use Committee (IACUC) with animal study protocol approval number TFBS2020-019 and Academia Sinica (approval number: 20-10-1526).

### SARS-CoV-2 S-2P protein antigen

Recombinant stabilized trimeric full length S protein expressed by stable CHO cell line was provided by Medigen Vaccine Biologics Corporation as described previously^28, 31^.

### Nanoemulsion Adjuvant and Vaccine Preparation

The 60% NE01 adjuvant was prepared by high shear homogenization of water, ethanol, cetylpyridinium chloride, non-ionic surfactants, and highly refined soybean oil to form an oil-in-water nanoemulsion with a mean particle size of ~400 nm as described previously^32^. The vaccine was prepared by mixing S-2P with NE01 adjuvant for a final concentration of 10 µg of S-2P with 20% NE01/dose.

### Immunization and challenge of hamsters

Hamsters were grouped into six groups A-F (n = 8 for Groups A to E, n = 6 for Group F) as shown in Table 1 and immunized with the following regimens:

**Table 1.**

| Groups | Amount of S-2P protein |  |  |
| --- | --- | --- | --- |
|  | First immunization | Second immunization | Third immunization |
| **A: IMx2 + INx1** | 3 µg | 3 µg (day 21) | 10 µg (day 105) |
| **B: IMx1 + INx1** | 3 µg | 10 µg (day 105) |  |
| **C: IMx2** | 3 µg | 3 µg (day 21) | - |
| **D: IMx1** | 3 µg | - | - |
| **E: DDIMx2 + INx1** | 0.75 µg | 0.75 µg (day 21) | 10 µg (day 105) |
| **F: Negative control** | - | - | - |

- Group A (IMx2 + INx1): Two intramuscular doses of 0.1 mL of MVC-COV1901 (3 μg of S-2P adjuvanted with 150 μg of CpG 1018 and 75 μg of aluminum hydroxide) on days 0 and 21 and followed by intranasal immunization with one dose of S-2P-NE01 vaccine (10 μg) on day 105.
- Group B (IMx1 + INx1): One intramuscular dose of 0.1 mL of MVC-COV1901 (3 μg of S-2P adjuvanted with 150 μg of CpG 1018 and 75 μg of aluminum hydroxide) on day 0 and intranasally with one dose of S-2P-NE01 (10 μg) on day 105.
- Group C (IMx2): Two intramuscular doses of 0.1 mL of MVC-COV1901 (3 μg of S-2P adjuvanted with 150 μg of CpG 1018 and 75 μg of aluminum hydroxide) on days 0 and 21.
- Group D (IMx1): One intramuscular dose of 0.1 mL of MVC-COV1901 (3 μg of S-2P adjuvanted with 150 μg of CpG 1018 and 75 μg of aluminum hydroxide) on day 0.
- Group E (DDIMx2 + INx1): Dose down (DD) group with two intramuscular doses of 25 μL of MVC-COV1901 (i.e. 0.75 μg of S-2P adjuvanted with 37.5 μg of CpG 1018 and 18.75 μg of aluminum hydroxide on days 0 and 21 followed by intranasal immunization with one dose of S-2P-NE01 vaccine (10 μg) on day 105
- Group F (NC): Unimmunized negative control.

Serum samples were harvested on day 91 and day 126 (i.e. 3 weeks after the third immunization) and sera derived from the bleeds were subjected to pseudovirus neutralization assay as in the following section.

### Pseudovirus neutralization assay

Hamster sera were analyzed for neutralizing antibody titers using pseudovirus composed of lentivirus expressing full-length wild type Wuhan-Hu-1 strain SARS-CoV-2 spike protein as described previously^28^. Briefly, sera were heat-inactivated, serially diluted 2-fold in MEM with 2% FBS and mixed with equal volumes of pseudovirus. The samples were incubated at 37°C for 1 hour before addition to plated HEK293-hACE2 cells. Cells were lysed 72 hours post incubation and relative luciferase units (RLU) were measured. ID50 and ID90 (50% and 90% inhibition dilution titers) were calculated deeming uninfected cells as 100% and virus transduced control as 0%.

### Hamster challenge with SARS-CoV-2

Hamsters were challenged with 1 × 10^4^ PFU of SARS-CoV-2 4-5 weeks after intranasal booster administration, as described previously^31^. Four hamsters from each group A-E and three hamsters from group F were then sacrificed three- and six-days post challenge for viral load and pathology in lungs along with collection of nasal wash for upper respiratory viral load. Body weight and survival for each hamster were recorded daily post challenge until sacrifice. Euthanasia, viral load and histopathological examination were performed as described earlier^31^.

### Quantification of viral titer by cell culture infectious assay (TCID50)

The viral titer determination from lung tissue was performed as described previously^31^. In brief, lungs were homogenized, clarified by centrifugation and supernatant was diluted 10-fold and plated onto Vero cells in quadruplicate for live virus estimation. Similarly for nasal wash, the sample was centrifuged, diluted, and plated onto Vero cells. Cells were fixed and stained, and TCID50/mL was calculated by the Reed and Muench method.

### Real-time PCR for SARS-CoV-2 RNA Quantification

SARS-CoV-2 RNA levels were measured using an established RT-PCR method to detect envelope gene of SARS-CoV-2 genome^33^. RNA obtained from both lungs and nasal wash was analyzed for SARS-CoV-2 RNA levels as described previously^31, 34^.

### Histopathology

As described previously^31, 35^, the left lungs of the hamsters were fixed with 4% paraformaldehyde for 1-week. The lungs were trimmed, processed, paraffin embedded, sectioned and stained with Hematoxylin and Eosin (H&E) followed by microscopic scoring. The scoring system was performed similar to previous experiments where nine different areas of the lung sections are scored individually and averaged^31^.

### Statistical analysis

The analysis package in Prism 6.01 (GraphPad) was used for statistical analysis. Spearman’s rank correlation coefficient and linear regression were calculated. Kruskal-Wallis with corrected Dunn’s multiple comparisons test and two-way ANOVA with Dunnett test for multiple comparison were used to calculate significance. * = p < 0.05, ** = p < 0.01, *** = p < 0.001, **** = p < 0.0001

## Results

To investigate the efficacy of an intranasally (IN) administered vaccine in animals previously vaccinated twice intramuscularly (IM) but likely experiencing a waning immune response over time, we devised an experimental plan where golden Syrian hamsters were challenged intranasally with SARS-CoV-2 virus after one or two standard IM vaccinations followed by one IN booster vaccination (Figure 1).

**Figure 1.**
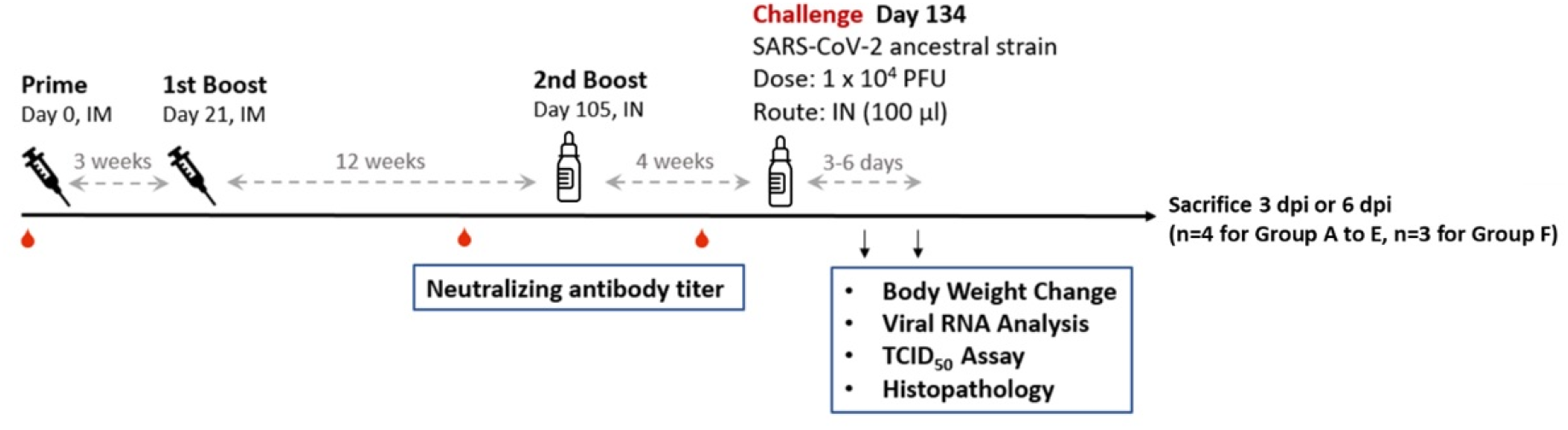
Design of the hamster challenge study. Hamsters (N=8 for group A-E and N=6 for group F) were immunized once (groups B and D), or twice (groups A, C and E) at three weeks apart for intramuscular immunization. Groups A, B and E were boosted by single intranasal immunization at the end of 12 weeks post last IM immunization. Serum samples were taken for immunogenicity assays at 91 days and 126 days after the first immunization. At 134 days after the first immunization, hamsters were challenged with 10^4^ PFU of SARS-CoV-2 ancestral strain. The animals were euthanized on the third or sixth day after infection for necropsy and tissue sampling to determine viral load. Body weight of individual hamster was tracked daily up to the time of sacrifice.

As a first analytical step, sera were examined in pseudovirus neutralization assays to assess levels of SARS-CoV-2 spike protein-specific antibody as an indicator of vaccine-induced immunity. The ID50 results for Day 91 prior to IN vaccination show that the second IM injection (Group A) significantly enhances immunogenicity compared to one dose of IM injection (Group B) alone, with ID50 GMTs of 1,460 and 272, respectively (Figure 2A). Importantly, immunogenicity is significantly enhanced by one dose of IN booster, with ID50 GMTs 3,395 and 4,119 in Groups A and B at Day 126, respectively. These results suggested that IN vaccination enhanced immunogenicity even when significant levels of immunogenicity have already been generated by IMx1 or IMx2. Surprisingly, two doses of IM injection of one-quarter amount of antigen and adjuvants (DDIMx2) followed by an IN vaccination induced a strong neutralizing antibody response with an ID50 GMT of 3,797, comparable to that of IMx1 or IMx2. As expected, IM vaccination without follow up with IN vaccination had reduction in neutralizing antibody titers by day 126, with the IMx2 group (Group C) having ID50 GMTs of 1,462 and 1,361 on days 91 and 126, respectively; IMx1 (Group D) had the lowest GMTs out of all vaccinated groups at 283 and 174 on days 91 and 126, respectively. Not surprisingly, data for the ID90 titers followed the same pattern (Figure 2B). First, the accuracy of measurements for vaccinated groups was not compromised by the upper assay limit. Second, the ID90 data reflected the relative difference of ID50 titers in groups A-F.

**Figure 2.**
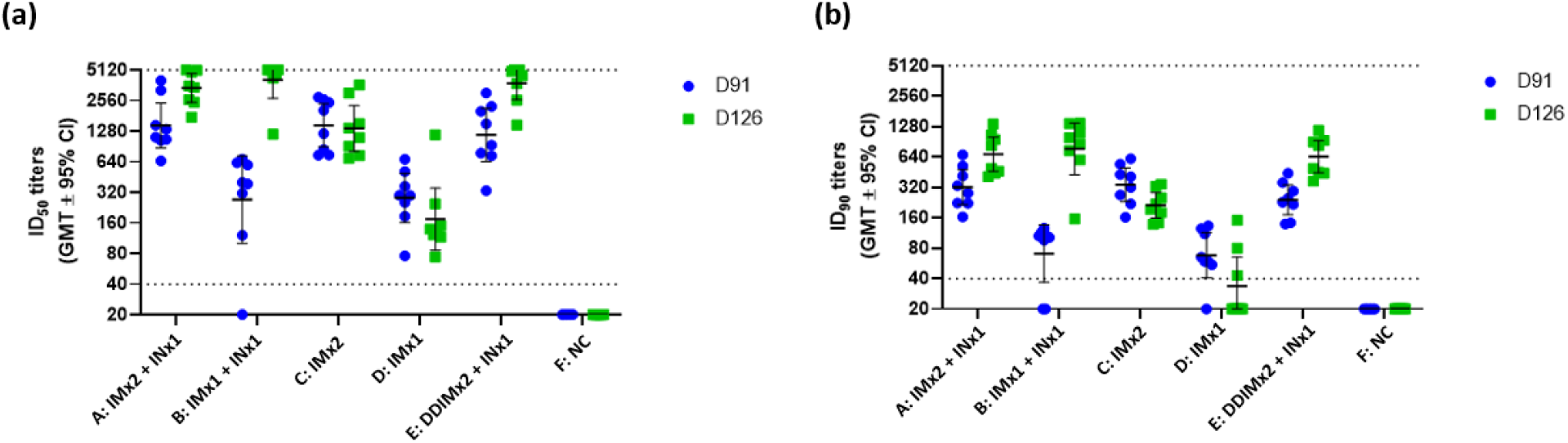
Induction of neutralizing antibodies in hamsters at 91 days and 126 days after first immunization. Hamsters (N=8 for group A-E and N=6 for group F) were immunized once (groups B and D), or twice (groups A, C and E) at three weeks apart for intramuscular immunization. Groups A, B and E were boosted by single intranasal immunization at the end of 12 weeks post last IM immunization. Serum samples were taken for immunogenicity assays at 91 days and 126 days after the first immunization. The antisera were subjected to neutralization assay with pseudovirus expressing SARS-CoV-2 spike protein to determine the ID50 (left) and ID90 (right) titers of neutralization antibodies. Each dot represents the serum sample neutralizing titer from each animal. Bars indicate geometric mean titers (GMT) and error bars indicate 95% confidence intervals. Dotted lines represent lower and upper limits of detection (40 and 5120, respectively, for both ID50 and ID90)

Based on body weight measurements after SARS-CoV-2 challenge (Figure 3), the immune response that was generated after vaccination appears to be at least partially protective against SARS-CoV-2 infection. Specifically, body weight decreased significantly by day 6 in the unvaccinated control group (Group F) after the viral challenge, while all other groups suffered little or no weight loss. Protective antibody generation appears to be robust enough even in the IMx1 animal group that achieved significantly lower antibody titers than all other vaccinated groups (compare Figures 2 and 3, Group D).

**Figure 3.**
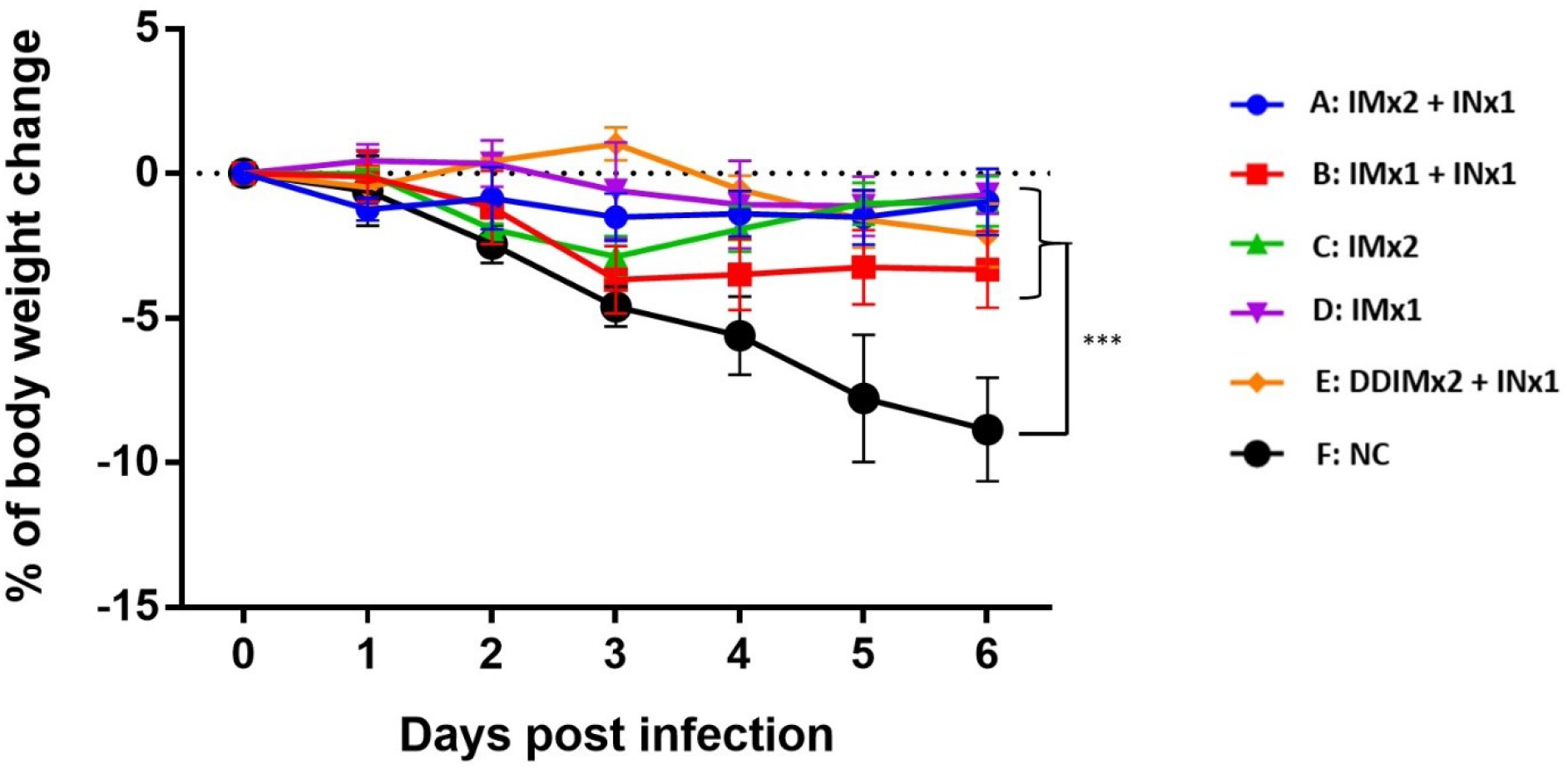
Changes in body weight post-SARS-CoV-2 infection. Results are presented as the mean ± S.E.M. Differences between negative control group and other experimental groups of animals were analyzed by two-way ANOVA; * p<0.05, ** p<0.01, ***p<0.001.

Intranasal vaccine booster efficacy was determined by measuring the viral load in lung and nasal wash by measuring viral titers using the TCID50 assay (Figure 4). Antibody levels suggested by the neutralization assays were clearly reflected by lung virus titers three days after viral challenge, in which all vaccinated groups had significantly reduced viral load compared to the control group (Figure 4a). Most importantly, the data showed a protective effect of the intranasal booster vaccination. Three days post challenge, viral titers were observed only in animals that did not receive the intranasal booster, while all groups that received IN vaccination had undetectable virus in lung homogenates (Figure 4a). Viral titers were reduced to undetectable levels on day 6 in the lungs of all hamsters except for the animals in the control group (Figure 4). In nasal wash, groups receiving only IM vaccination (Groups C and D) had no significantly different viral titers than the control group at 3 days post infection (Figure 4b). At the end of three days post challenge, animals that received the intranasal booster showed undetectable virus in nasal washes with the exception of the group which received the reduced dose of IM vaccination (Group E), although it was significantly lower than control. However, by day 6, virus was cleared from all the groups including the control (Figure 4b). Viral load was also measured by viral RNA titer (Figure 5). Lung RNA titers were significantly lower in the IN boosted groups than the control group on day 3 (Figure 5a). Similarly, in the nasal wash, RNA levels were significantly lower than in the IN boosted groups than in the control group on day 3, while on day 6 levels of viral RNA in all vaccinated groups were significantly reduced compared to the control even though these levels in all groups remained high (>10^7^ μg) (Figure 5b).

**Figure 4.**
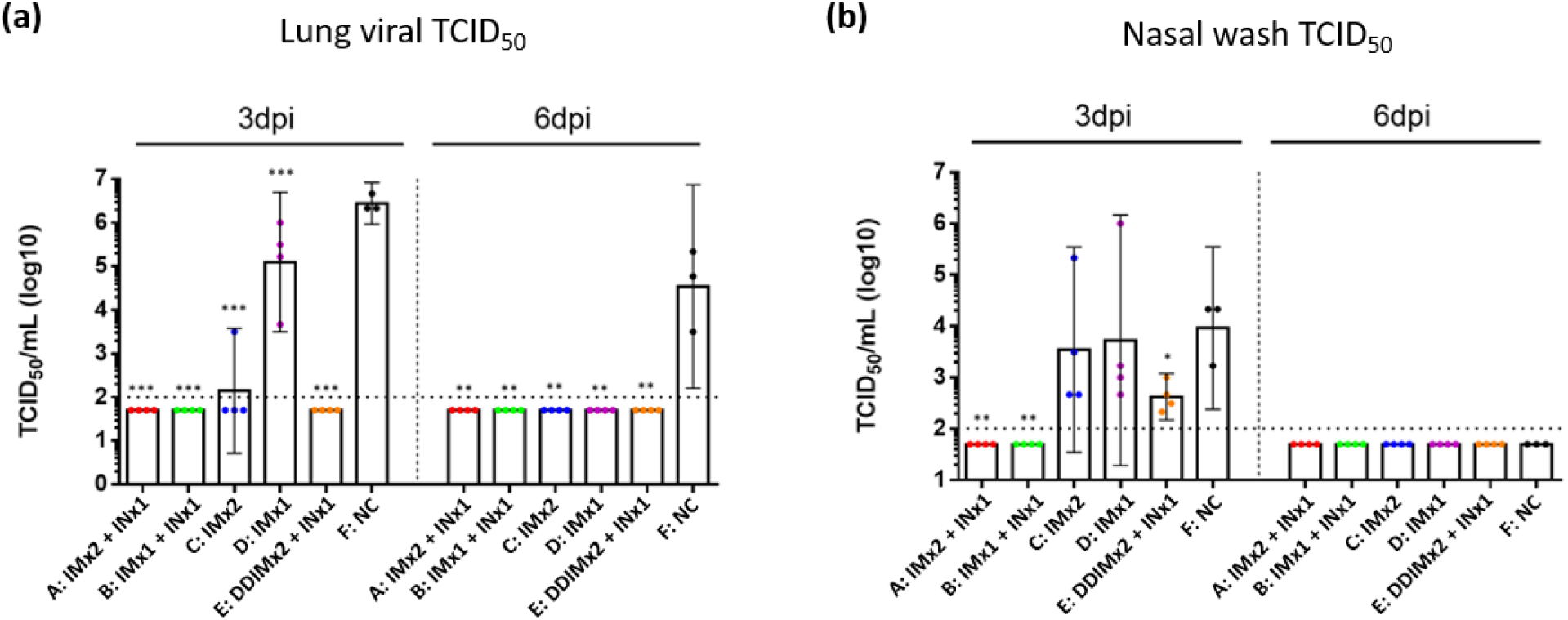
Tissue culture infectious dose (TCID50) from (A) lung homogenates; (B) nasal wash of hamsters at day 3 and 6 post-SARS-CoV-2 infections. Results are presented as the geometric mean values. Differences between negative control group and other experimental groups of animals were analyzed by student’s t test; * p<0.05, ** p<0.01, ***p<0.001. Dotted line: limit of detection.

**Figure 5.**
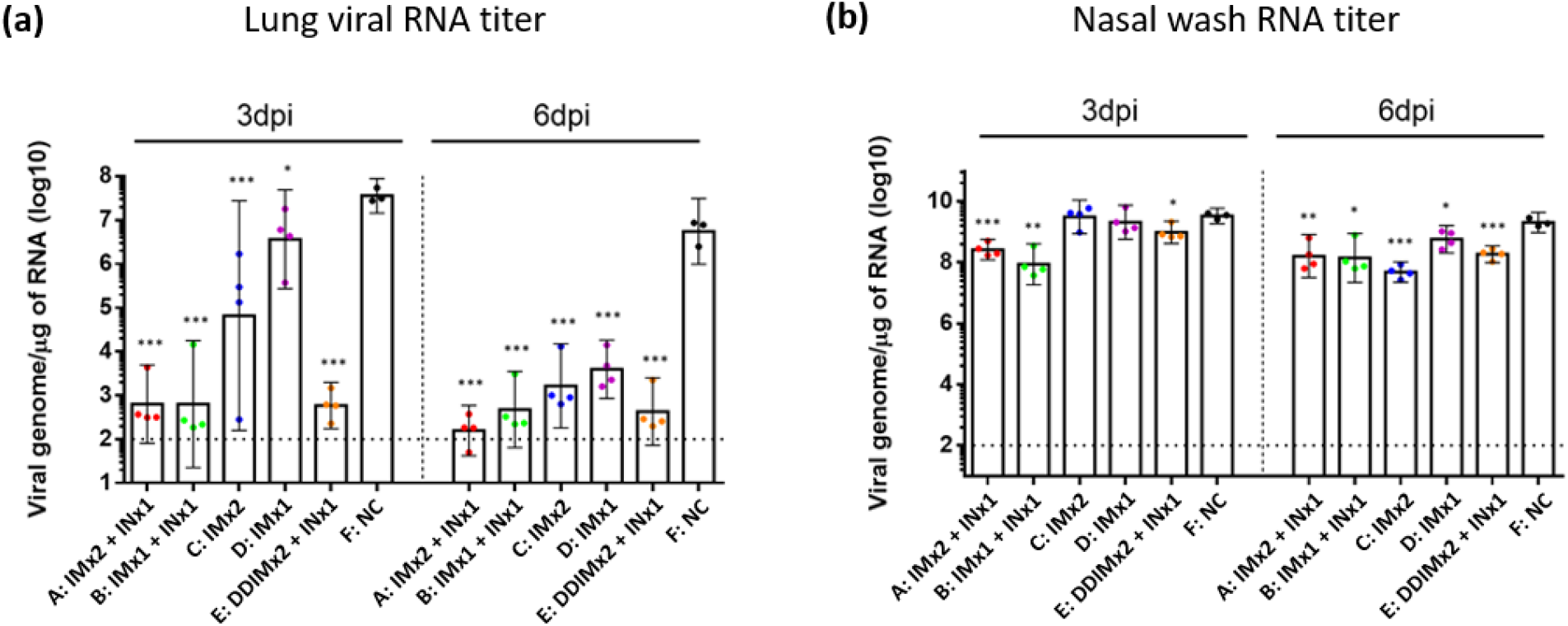
Viral RNA titer of hamster at days 3 and 6 post-SARS-CoV-2 infection. (A) lung viral RNA titer; (B) nasal wash at days 3 and 6 post-SARS-CoV-2 infection. Results are presented as the geometric mean values. Differences between negative control group and other experimental groups of animals were analyzed by student’s t test; * p<0.05, ** p<0.01, ***p<0.001. Dotted line: limit of detection.

Lung health was assessed by pathology scores to determine if intranasal vaccination can protect lung function (Figures 6 and S1). As shown in Figure 6, three days after viral challenge, lung pathology scores were intermediate between 2 and 3, and indistinguishable from the NC control for all treatment groups except group A (intramuscularly vaccinated twice and intranasally boosted). Only group A animals showed a statistically significant reduction in pathology score (p<0.05) compared to the non-treated NC control group, thus supporting that IN vaccination induces immune protection that achieves improved lung scores at least until Day 3. On Day 6, lung scores in the non-vaccinated NC control group had worsened from a score of 2.5 on Day 3 to a score of 4.5 with significant infiltration and necrosis (Figure S1), suggesting progression of SARS-CoV-2 in unvaccinated animals between Day 3 and Day 6. In contrast, pathology scores in all vaccinated animal groups displayed lung scores that were similar on Day 3 and Day 6.

**Figure 6.**
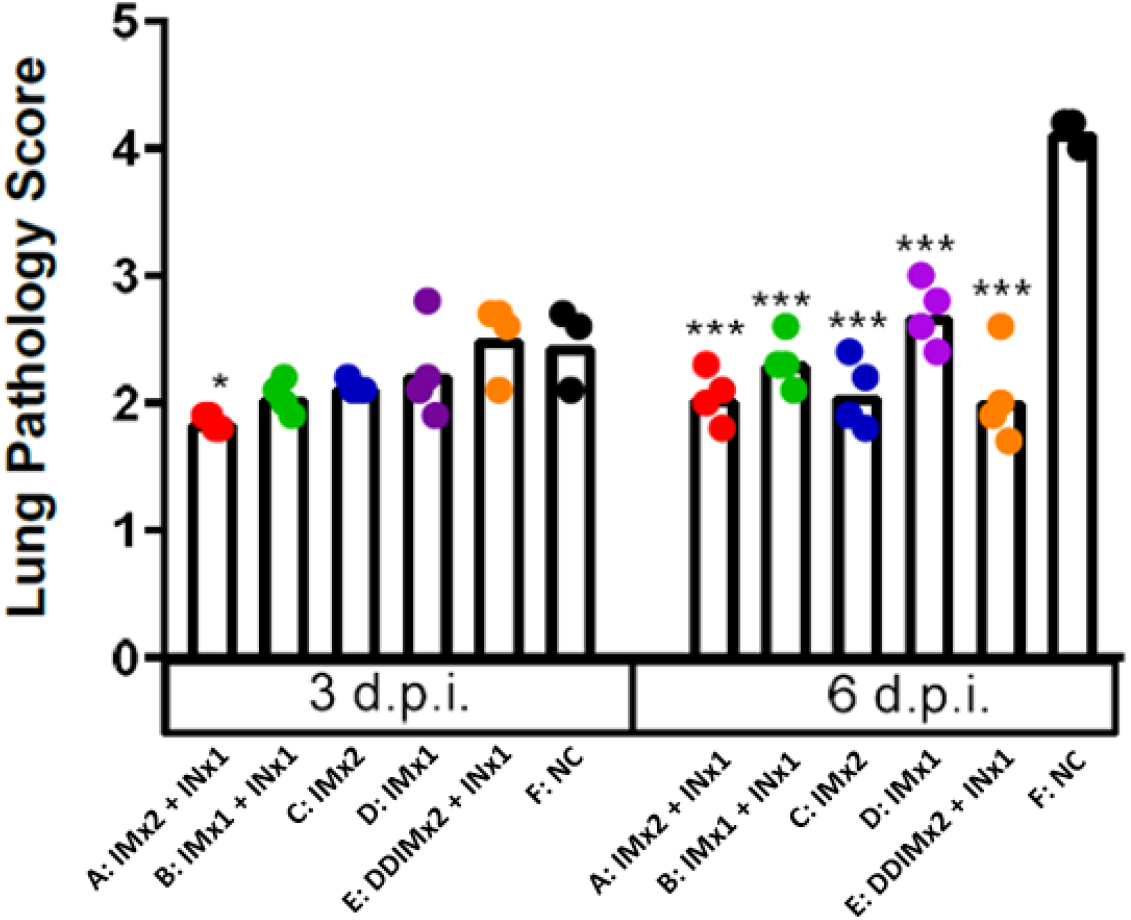
The clinical scoring of hamsters at day 3 and 6 post-SARS-CoV-2 infection. Results are presented as the mean values. Differences between negative control group and other experimental groups of animals were analyzed by student’s t test; * p<0.05, ** p<0.01, ***p<0.001.

## Discussion

In this study, we have investigated the booster effect of a single intranasal vaccination as a booster after two doses of intramuscular vaccine in hamsters to address the waning immunity against SARS-CoV-2 and inadequate protection from viral carriage following intramuscular vaccination. Here, the efficacy of the intranasal vaccine as a booster was demonstrated in the hamster challenge model where two intramuscular doses of MVC-COV1901 vaccine followed by a single dose of intranasal NE01-adjuvanted vaccine induced protection from intranasal challenge of SARS-CoV-2. Similar levels of neutralizing antibody titers were seen in all IN boosted groups on day 126, implying that IN vaccination can boost systemic immunity regardless of previous vaccination (Figure 2). All IN boosted groups also showed significantly reduced virus titers and viral RNA levels at 3 days after virus challenge (Figures 4 and 5). Although viral transmission assay was not performed in this study, IN vaccination could inhibit viral transmission based on the fact that virus was undetectable in the nasal washes of most of the IN vaccinated animals.

The data of this initial study in a hamster model suggested that IN vaccination does safely generate immunogenicity towards SARS-CoV-2. Specifically, the IN vaccine strongly boosted serum antibody titer by Day 126 even after only one IM immunization (Figure 2). Thus, the IN vaccine may have potential utility as a booster shot after immunity from IM vaccination(s) has declined. Importantly, for the key objectives of this study, and consistent with the serum antibody assays, quantitation of infectious virus in lung tissue and nasal washes three days after viral challenge showed that single intranasal booster can eliminate detectable virus after one previous IM vaccination dose (Figures 4 and 5). Finally, body weight measurements and lung pathology scores showed significant weight loss and high pathology scores only for the unvaccinated control group (Figures 3 and 6). We conclude that IN vaccination with our vaccine formulation demonstrated no significant safety concerns. Overall, this data set warrants further investigation of this intranasal vaccine. We are also encouraged that the vaccine generated noticeable benefit as a booster, suggesting that the vaccine may also be effective as a primary course of vaccination. In the IN ChAdOx1 nCoV-19 animal study, the investigators found that two IN doses of ChAdOx1 nCoV-19 in macaques and hamsters were able to induce robust IgA and IgG responses and reduced viral shedding from the upper respiratory tract as well as lowered viral load in lower respiratory tract^27^. In the same study, intranasal vaccination also protected hamsters from transmission of the virus when vaccinated hamsters were co-housed with infected hamsters^29^.

The intranasally delivered vaccine of SARS-CoV-2 S-2P antigen used in this study was formulated with nanoemulsion adjuvant NE01. In animal models (Respiratory Syncytial Virus in cotton rats and pandemic flu in ferrets), intranasal vaccines adjuvanted with NE01 elicited mucosal and systemic immunity that prevented both disease and nasal colonization^36, 37^. Additionally, homing of memory cells and induction of mucosal immunity in distant mucosal tissues were also achieved in the above studies. The safety profile of NE01 and its stability (at 5°C) that allows for normal storage and handling, together with ease of administration that does not require needles or highly trained personnel, makes NE01-formulated vaccines attractive, especially for low-income/low infrastructure countries^38^.

From a practical point of view, disease control is also complicated by vaccine hesitancy. From a biological point of view, the currently licensed vaccines are administered intramuscularly which means that immune protection against virus is difficult to achieve in the primary and secondary sites of infection, i.e. the nasal and upper respiratory tract passages, respectively^26^. For these reasons, intranasal administration of the vaccine formulated as a convenient nasal spray might offer an attractive solution to both problems.

Admittedly, the current study has limitations. Specifically, we did not measure the levels of IgA as an indicator of mucosal immunity in the nasal, or lung tissues, and we did not investigate the homing of T- and B-cells to mucosal tissues. We also did not test whether using IN as a primary series of vaccination (as in the previously referred IN ChAdOx1 nCoV-19 study) can induce the same level of immunity and protection as seen when using IN as a booster. Detection of subgenomic RNA should be done in the future studies to better corroborate the TCID50 live virus count with viral RNA titer as genomic RNA can detect RNA from both live and dead virus, as well as viral RNAs released from dead cells, whereas subgenomic RNAs are only found in actively replicating cells but not packaged in virions^39^.

The original Wuhan and other variants of SARS-CoV-2 have been replaced by the circulating Omicron and Delta variants^9^ and most likely the virus will continue to evolve and produce new VoCs. The outcomes of IN vaccination with MVC-COV1901 adjuvanted with NE01 against VoCs was not investigated in this study. However, our previous data have shown that administration of booster dose of MVC-COV1901 of either wildtype S-2P or Beta variant of S-2P could confer protection against Delta variant challenge in hamsters^40^. In addition, three doses of MVC-COV1901 also improved immunogenicity against VoCs compared to two doses of MVC-COV1901 in a clinical trial^41^. Therefore, it is reasonable to assume that the regimen of IM vaccination followed by IN booster will generate sufficiently broad protective immunity and protection against VoCs. The IN boosting that not only boosts systemic immunity but also induces mucosal immunity might be a solution to the current and likely persistent VoC problem.

## Acknowledgements

We are grateful for the participation of Dr. Han van den Bosch for manuscript review and constructive comments. We also thank team members at TFBS Bioscience Incorporation for hamster housing and immunization process. We thank the Biomedical Translation Research Center, Academia Sinica, Taiwan, for performing hamster challenge. In addition, we would like to acknowledge Dr. Yu-Chi Chou and his team at the RNAi Core Facility, Academia Sinica for the pseudovirus neutralization assay.

## Author Contributions

T.-Y. K. produced the S-2P antigens used in the study. C.-E. L., Y.-J. L., M.-Y. L., Y.-S. C., and C. C., S. G., A.F., and V. B. designed the study and experiments. Y.-J. L. and Y.-S. C. supervised the experiments at TFBS Bioscience and Academia Sinica. Y.-J. L., M.-Y.-L., Y.-S. C., and L. T.- C. L. analyzed the results. M.-Y. L., Y.-S. C., and L. T.-C. L. drafted the manuscript. All authors reviewed and approved of the final version of the manuscript.

## Figures

**Figure S1.**
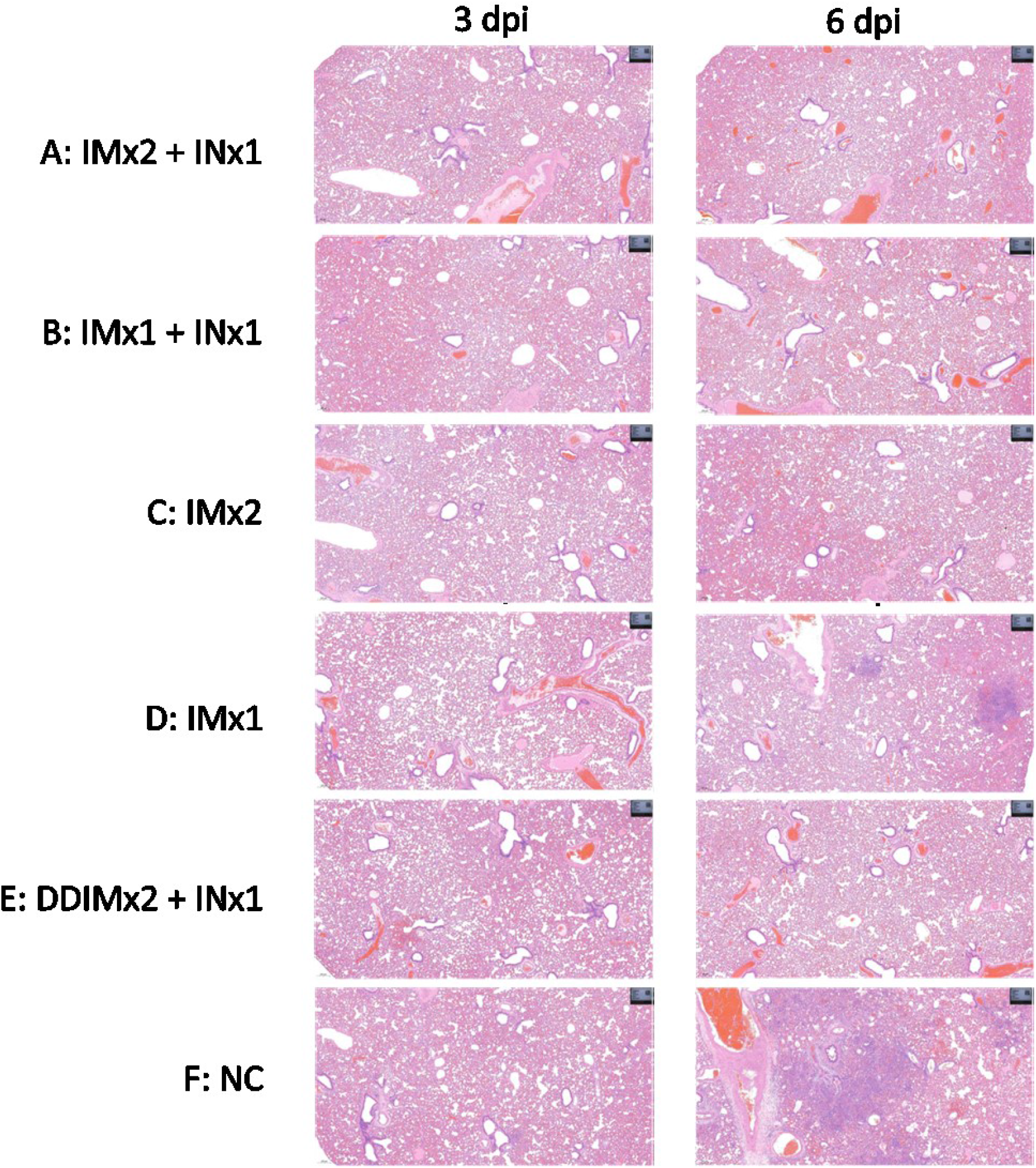
Representative histopathology sections of infected hamsters from Groups A to F at 3 d.p.i. or 6 d.p.i. The left lungs of hamsters were isolated and fixed in 4% paraformaldehyde for one week, sectioned and stained with Hematoxylin and Eosin for visualization.

